# A robust synthetic biology toolkit to advance carboxysome study and redesign

**DOI:** 10.1101/2024.10.08.617227

**Authors:** Daniel S. Trettel, Y Hoang, Anthony G. Vecchiarelli, Cesar R. Gonzalez-Esquer

## Abstract

Carboxysomes are polyhedral protein organelles that microorganisms use to facilitate carbon dioxide assimilation. They are composed of a modular protein shell which envelops an enzymatic core mainly comprised of physically coupled Rubisco and carbonic anhydrase. While the modular construction principles of carboxysomes make them attractive targets as customizable metabolic platforms, their size and complexity can be a hinderance. In this work, we design and validate a plasmid set – the pXpressome toolkit -in which α-carboxysomes are robustly expressed and remain intact and functional after purification. We tested this toolkit by introducing mutations which influence carboxysome structure and performance. We find that deletion of vertex-capping genes results in formation of larger carboxysomes while deletion of facet forming genes produces smaller particles, suggesting that adjusting the ratio of these proteins can rationally affect morphology. Through a series of fluorescently labeled constructs, we observe this toolkit leads to more uniform expression and better cell health than previously published carboxysome expression systems. Overall, the pXpressome toolkit facilitates the study and redesign of carboxysomes with robust performance and improved phenotype uniformity. The pXpressome toolkit will support efforts to remodel carboxysomes for enhanced carbon fixation or serve as a platform for other nanoencapsulation goals.

## Introduction

Bacteria have been increasingly found to organize and exert spatial control over their internal cytoplasm.^1,2^ Several instances have been described of bacteria using protein-bounded organelles that serve various cellular functions. Some, like encapsulins, can serve storage and biosynthetic roles.^3,4^ Others, like gas vesicles, allow host bacteria to respond to their environment and regulate their buoyancy.^5,6^ These partitions and their modular construction principles can be adapted for synthetic nanoencapsulation goals. Among these, bacterial microcompartments (BMCs) in particular serve as the predominant archetype of a modular bacterial organelle.^7^

BMCs play important roles in bacterial metabolism and fall into two subcategories depending on if they display catabolic or anabolic functions. Catabolic BMCs are called metabolosomes, a functionally diverse class of compartments,^8^ while anabolic BMCs are typified solely by the carbon-fixing carboxysomes. All BMCs consist of two distinct domains: a structurally conserved semipermeable protein shell^9-12^ and the enzymatic cargo encased within.^13^ These polyhedral structures are naturally over 100 nm in diameter and over 200 MDa in mass.^14,15^ It should be noted that synthetic derivatives are typically much smaller at ≤40 nm in diameter.^16-19^ The shells are made from 3 classes of structural proteins; hexamers (BMC-H), trimers (BMC-T) and pentamers (BMC-P).^7,8^ BMC-H/T compose the facets of the polyhedra while BMC-P cap the vertices.^20,21^ In contrast to other protein organelles like encapsulins, BMC display morphological and functional adaptability owing to their modular design principles.^22,23^

BMCs present unique engineering opportunities and challenges that stem from their modularity, size, and complexity. Despite the structural features of BMCs being well established, the size and genetic complexity of native systems can still hamper research efforts. For instance, the model α-carboxysome from *Halothiobacillus neapolitanus* encodes for 10 open reading frames (with one being translated into two different forms)^24^ across two distinct genomic loci that total to approximately 9 kilobases of DNA. The genetic structure of β-carboxysomes is even more convoluted and spread across numerous satellite loci^8^ where the specific expression parameters are only beginning to be understood.^25-27^ Metabolosomes face a similar problem where some, such as the 1,2-propanediol utilization BMC, can reach upwards of 20 kilobases in length and encode for nearly 20 unique proteins.^28^ Other metabolosomes, like ethanolamine microcompartments, may be encoded by a single locus but are subject to complex native riboregulation schemes.^29,30^ Structurally, BMCs are known to include thousands of individual proteins with copy numbers ranging from the tens to thousands of units.^14,31^ Entirely synthetic model systems exist but are usually under 40 nm in diameter and rely on non-native packaging approaches.^32-34^ Synthetic reconstruction of native-like BMCs is possible but can be tedious and commonly produce non-polyhedral assemblies^35,36^ likely due to an improper understanding and recapitulation of component stoichiometry and assembly kinetics.^37-39^

Classical strategies to manipulate BMCs have relied on genetic manipulation of native organisms which requires niche tools specific to ones’ particular system of study. Newer strategies have used recombination approaches to produce bacterial artificial chromosomes (BACs) which can produce functional metabolosomes from native induction.^40,41^ However, the size of these BACs limits the utility of common and established molecular biology approaches like PCR mutagenesis and Gibson assembly. Plasmid-based approaches would be ideal due to the genetic tractability of commonly used bacteria as well as their ability to be rapidly and accurately mutated and sequence validated. Several plasmid-based expression systems for BMC systems have been described in recent years for both metabolosomes^33^ and carboxysomes.^14,42-44^ These systems can drive expression of structurally and functionally intact BMCs and allow for their manipulation and study.

In this study, we introduce a new suite of plasmids for the heterologous expression and engineering of *H. neapolitanus* α-carboxysomes which we make available for widescale adoption. We focus on α-carboxysomes due to their important role in biological carbon fixation and thus the global carbon cycle. This plasmid toolkit, called the pXpressome system, offers simple and robust methods to enrich, test and modify α-carboxysomes for new or enhanced functions. We first show that the pXpressome system can produce structurally and functional intact carboxysomes which enables their characterization. We produce several mutants which reveal that the ratio of shell constituents (BMC-H and BMC-P) can control particle morphology. We next fuse a series of fluorescent proteins, spanning the entire visible light spectrum, to Rubisco to enable tracking and imaging of carboxysomes *in vivo*. Lastly, we show that the pXpressome system offers more consistent growth and expression phenotypes compared to other plasmid-based systems, likely owing to a more tightly regulated promoter. Overall, we showcase the utility of a robust plasmid toolkit that enables any researcher to design their own carbon-fixing nanoreactor.

## Results

### A system for the recombinant expression of α-carboxysomes

We chose to work with the model α-carboxysome from the chemoautotrophic bacteria *H. neapolitanus*. This α-carboxysome is natively made of Rubisco (CbbLS) and carbonic anhydrase (CsoSCA) coordinated by the full (CsoS2B) and short forms (csoS2A) of the disordered scaffold protein CsoS2 and encased within a protein shell (Figure 1A). The facets of the shell are made from three BMC-H proteins (CsoS1C, S1A, and S1B) while the vertices are capped by two BMC-P proteins (CsoS4A, S4B). The pXpressome plasmid encodes for all these components under control of the araBAD promoter. Our system differs from others by the promoters used (lacUV5 vs araBAD)^42^ and components encoded (Figure 1B). We opted for an araBAD promoter due to the higher degree of regulation control. We also only encoded the genes within the main carboxysome locus (Figure 1B), and thus excluded the BMC-T protein CsoS1D and the activases CbbOQ used by other methods^14^ to create a simplified format for manipulation.

**Figure 1:**
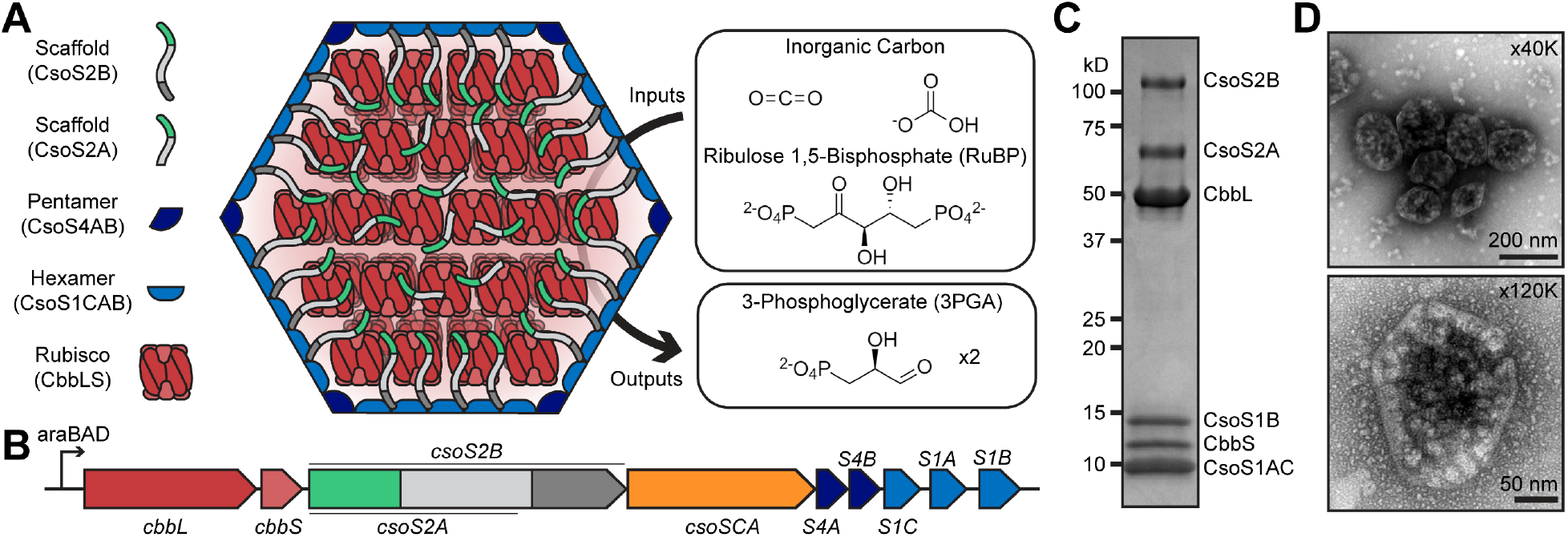
Production of intact recombinant α-carboxysomes. **(A)** The carboxysome is a multicomponent structure of Rubisco latticed within a polyhedral shell. It functions to combine inorganic carbon with RuBP to produce two molecules of 3PGA. Scaffolds anchor Rubisco subunits to the shell while also non-covalently crosslinking the interior lattice. Carbonic anhydrase, which is known to associate with Rubisco, is omitted for clarity. **(B)** A schematic of the minimal *H. neapolitanus* α-carboxysome operon used in this study. Expression is driven by the araBAD promoter and accessory components, like CsoS1D and CbbOQ, are not included in the design. **(C)** An SDS-PAGE profile of α-carboxysomes enriched from TOP10 *E. coli*. Bands correspond to the known protein components of the carboxysome with minimal contaminating components. **(D)** Carboxysome samples were negative stained and imaged with transmission electron microscopy, confirming appropriate morphology. Distinct bounded structures between 100-200 nm diameter are observed. Magnification values are labeled in the top right corners.

We used a common lab strain of *E. coli* (TOP10) as a host for transformation and recombinant production of carboxysomes. This is due to TOP10 *E. coli* being unable to metabolize the arabinose needed for induction which otherwise limits carboxysome expression in other common strains like DH5α (Figure S1). Samples produce a banding profile on SDS-PAGE like other work in natively produced *H. neapolitanus* carboxysomes and matches the anticipated masses of the components^14,42^ with minimal background (Figure 1C). Samples were then evaluated for correct particle morphology by transmission electron microscopy (TEM) which confirmed the presence of assembled α-carboxysomes (Figure 1D).

### Manipulation and analysis of recombinant carboxysome morphology and activity

One benefit of a plasmid expression platform is the ability to rapidly generate, screen, and study mutant designs. To exemplify this point, we produced knockout mutants of the carbonic anhydrase (ΔCsoSCA), a pentamer (ΔCsoS4A) and a hexamer (ΔCsoS1C) using standard PCR-based mutagenesis approaches. All mutants were able to produce carboxysome-like particles with similar banding profiles and purity on SDS-PAGE compared to the wild type. An exception is ΔCsoS1C which had a visibly reduced amount of CsoS2A (Figure 2A), the short form of CsoS2 known to non-covalently crosslink Rubisco^45^ while lacking the C-terminal domain responsible for shell binding.^46,47^ TEM analysis again revealed the presence of intact particles from all samples (Figure 2B). One curiosity is that the ΔCsoS4A sample produced a small population of elongated carboxysomes, similar to those reported elsewhere for complete BMC-P deletions,^48-50^ and recently reported nanocones^36^ (Figure S2). Dynamic light scattering (DLS) analysis revealed that wild type and ΔCsoSCA samples have similar diameters (given by the Z-average) of ∼150 nm as expected since the carbonic anhydrase does not play a significant structural role (Figure 2C). In contrast, ΔCsoS4A samples were larger (∼170 nm) and ΔCsoS1C particles were smaller (∼120 nm) than the wild type which suggests that the abundancies of these structural components can fine tune morphology.

**Figure 2:**
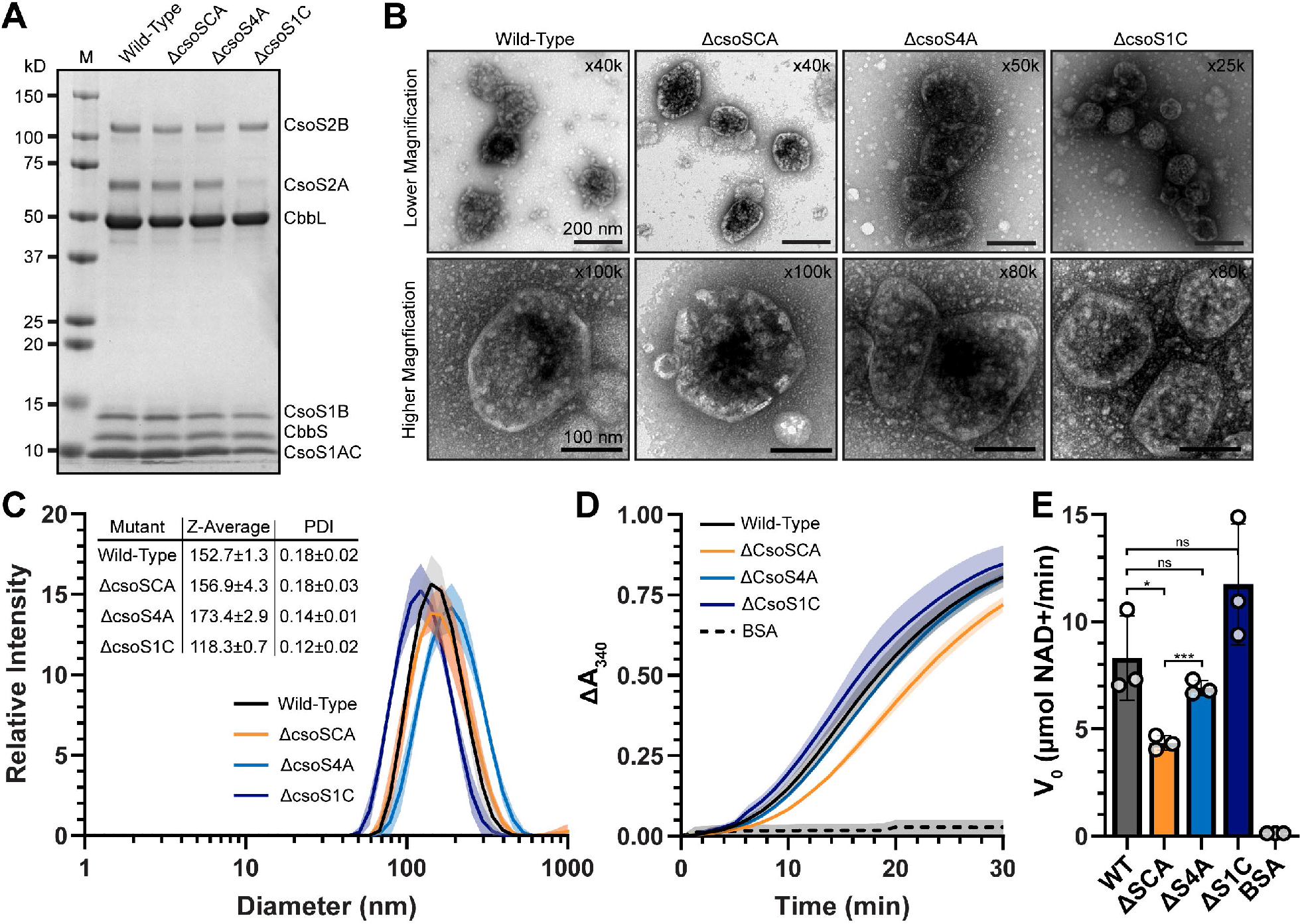
Manipulation and analysis of carboxysomes. **(A)** SDS-PAGE profile of several carboxysome knockouts that were produced and enriched from E. coli. Banding patterns are mostly similar, except ΔCsoS1C has a noticeably lower amount of CsoS2A. The gel was performed with 6 µg of each sample. **(B)** Negative stain transmission electron microscopy analysis of mutant carboxysomes. All mutants can form intact particles. Magnification levels are given in the top right corners. **(C)** Dynamic light scattering analysis reveals that mutant carboxysomes can differ in hydrodynamic radius (Z-average) with the CsoS4A (BMC-P) knockout giving larger particles and CsoS1C knockout (BMC-H) giving smaller particles. **(D)** A simple activity assay shows that all carboxysome mutants are enzymatically active, albeit giving different V_0_ **(E)** under these specific conditions.

SDS-PAGE, TEM and DLS evidence shows that our system can produce a range of structurally intact mutant carboxysomes. While these particles are all structurally intact, this does not mean that they are functional, which is critical for adaptation and screening efforts. To test this, we subjected carboxysome samples to a coupled-enzyme assay which takes the output 3-phosphoglycerate (3PGA) to oxidize NADH into NAD+, thereby providing a spectroscopic output by the decrease of absorbance at 340 nm for activity detection.^51^ This spectroscopic assay shows that all mutants are enzymatically active to varying degrees (Figure 2D). In contrast, a bovine serum albumin (BSA) negative control was unable to produce activity, which shows the assays specificity. Initial velocity (V_0_, in simple terms of µmol NAD+ produced per minute) was assessed and showed that, as expected, the carbonic anhydrase-lacking mutants have the slowest activity (Figure 2E). All other mutants were not significantly different from the wild type samples. Still, these results show that carboxysomes are compositionally plastic and that very plasticity influences their morphology.

### Development of a carboxysome palette for cytological experiments

The data presented so far show that the pXpressome system can produce structurally and functionally intact carboxysomes for *in vitro* analysis. We sought to increase the utility of these system by producing fluorescently tagged carboxysomes for *in vivo* studies as well. We chose to fuse various fluorescent proteins to the C-terminus of the small Rubisco subunit CbbS as an established tagged site, although recent evidence suggests that the N-terminus of CsoSCA can also work as an inclusion tag.^52^ Our palette includes, in increasing spectral order, (1) mTagBFP2, (2) mJuniper, (3) mChartreuse, (4) mNeonGreen, (5) mLemon, (6) mScarlet-I3 and (7) mLychee. Fluorescent proteins were chosen based on available evidence of their monomeric oligomer state, brightness, optical stability, fast maturation speeds and, in some cases, their utility in super-resolution imaging methods (Preprint).^53-56^

Control constructs expressing select fluorescent proteins (mTagBFP2, mNeonGreen and mScarlet-I3) from the araBAD promoter all show a uniform cellular fluorescence distribution within cells after overnight induction (Figure 3A, top). Overnight expression of these fluorophores fused to the C-terminus of CbbS, in contrast, reveal a distinct distribution pattern (Figure 3A, bottom) suggesting specific packaging of the fluorophore within carboxysomes. These overnight induced cultures show that fluorescence may not be entirely cytosolic but instead pushed towards ends of the cells and exclusion from the nucleoid. To test this, we induced carboxysome expression in cells at mid-log phase and imaged carboxysome expression at several early stages while staining the nucleoid with SYTO9. Carboxysome expression is apparent after 90 minutes of induction, as shown by the presence of red foci (Figure 3B, top). These foci were at the poles of cells and did not overlap with the stained nucleoid. This effect is even more pronounced after 4.5 hours of induction (Figure 3B, bottom). These data suggest that recombinant carboxysomes cluster at the cell poles due to nucleoid exclusion (Figure 3B, right). This contrasts with native systems that also encode the McdAB carboxysome distribution proteins which is not encoded in our pXpressome system.^57,58^ Our TEM evidence shows that this does not influence final morphology of the particles and they likely nucleate, mature and exist largely excluded from the nucleoid and densely clustered at the poles (Figure 3B, right). The ability to fluorescently tag CbbS allowed us to expand the palette of fluorescent fusions to include a total of 7 designs with excitation and emission profiles covering the visible light spectrum (Figure 3C). This palette expands utilities beyond any one fluorophore to accommodate any set of lasers and/or filters. Further, this palette can be used in combination with other fluorescent moieties like nucleic acid or membrane stains to track carboxysome formation and behavior.

**Figure 3:**
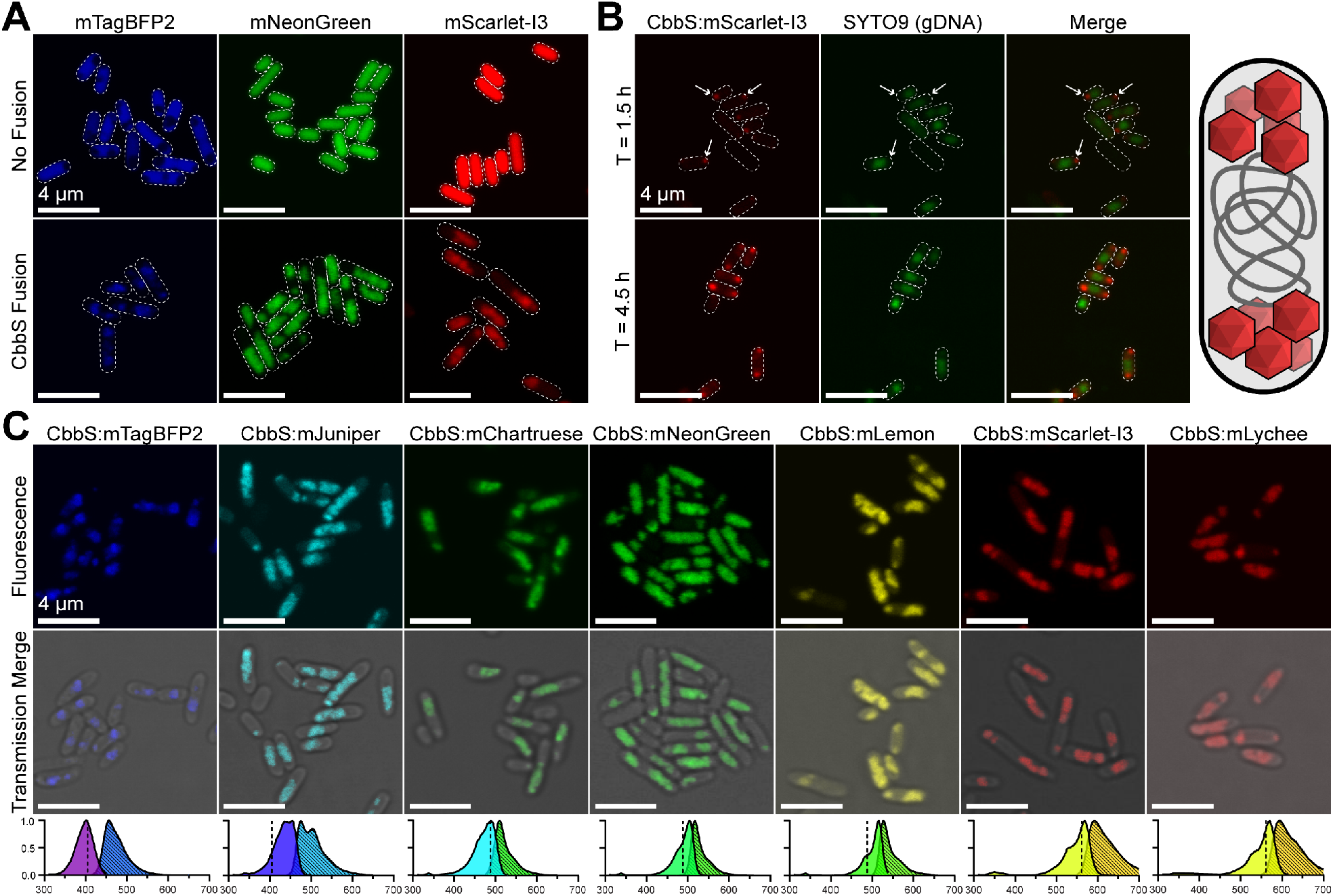
Developing a fluorescent carboxysome palette. **(A)** CbbS was C-terminally fused to various fluorescent proteins. Compared to these fluorescent proteins expressed alone, the CbbS fusions produce a unique organizational pattern within cells in agreeance with multiple particle formation. **(B)** Imaging early timepoints of carboxysome production reveals that recombinant carboxysomes are likely genome excluded. Carboxysomes tagged with mScarlet-I3 initially form at the cell poles and are excluded from gDNA (stained with SYTO9) which is more central. **(C)** A carboxysome palette was designed to meet researchers’ capabilities and needs across the visible light spectrum. The excitation and emission profiles (dashed fill) of each fluorophore are shown below each sample with the excitation laser lines used for imaging being represented by a dashed line. The spectral properties for each fluorescent protein came from FPbase and were plotted in GraphPad Prism.

### The pXpressome System Leads to Robust Expression and Uniform Phenotypes

Other publicly available plasmids exist that express *H. neapolitanus* α-carboxysomes. One such instance, called pHnCB, uses the lacUV5 promoter to successfully drive expression of carboxysomes^42^ which differs from our systems’ use of the araBAD promoter. We sought to test if these differences had functional consequences for expression and cell health. We tested this by comparing the expression of α-carboxysomes from pHnCB using a mTurquoise2 (mTQ2)^59^ reporter C-terminally fused to CbbS and compared it to our similarly designed pXpressome palette. All designs can drive expression of their respective fluorescent reporters (Figure 4A). However, apparent morphological differences between samples motivated us to quantify the effects of synthetic carboxysome expression on cell morphology. Cells expressing carboxysomes from pHnCB were elongated and slightly wider compared to pXpressome carrying cells (Figure 4B and 4C). Specifically, pHnCB expressing cells had a median length and width of 5.8 and 1.4 µm, respectively, while all pXpressome expressing cells were smaller with median widths between 3.2 – 3.5 µm and widths between 1.1 – 1.2 µm. Further, cells expressing from pXpressome showed morphological homogeneity across all designs while pHnCB cells showed far greater variance. These results may indicate leaky expression which negatively impacts cell division and/or cell health.

**Figure 4:**
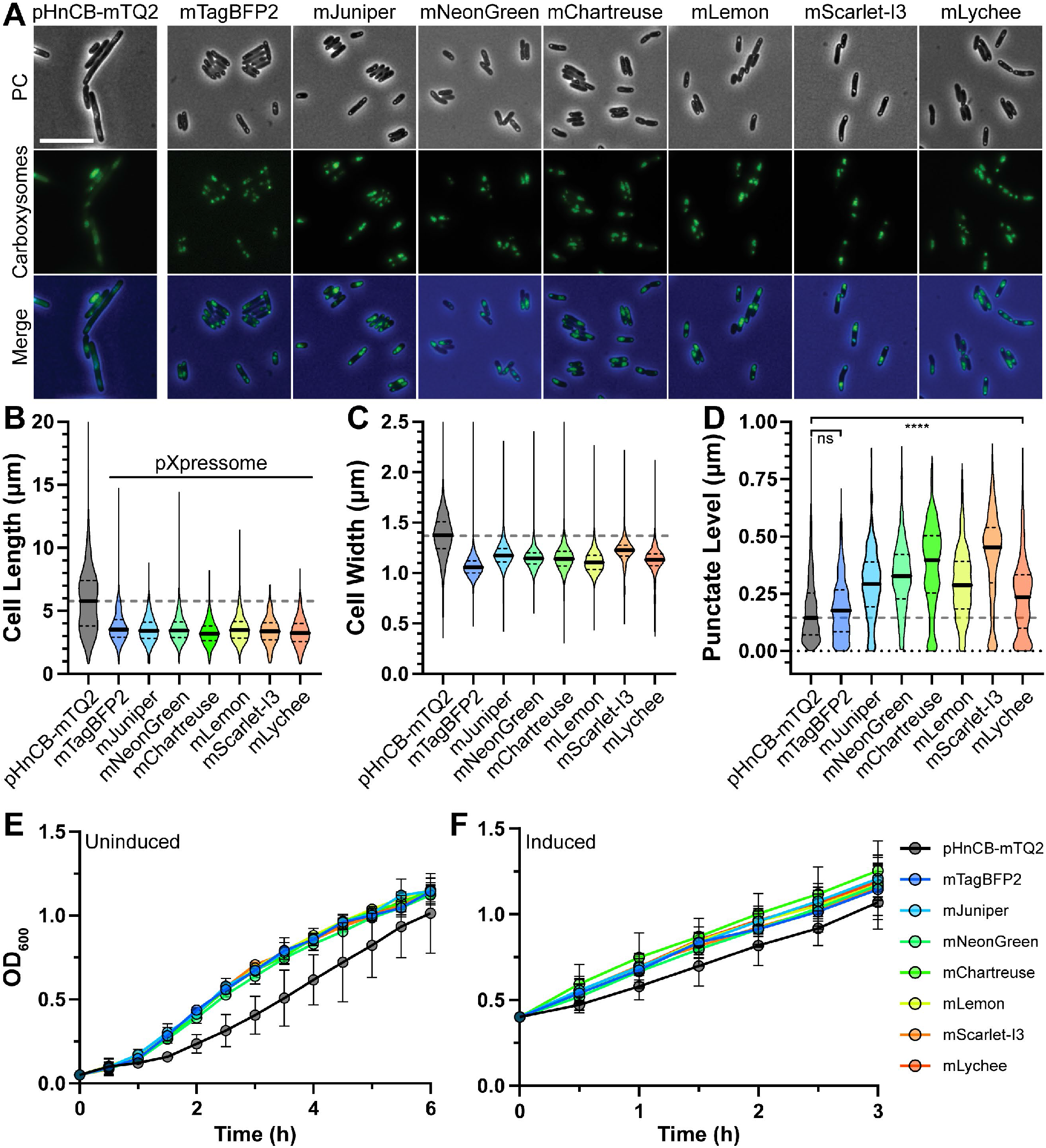
Analysis of phenotype consistency across plasmid designs. **(A)** Wide field fluorescence and phase-contrast microscopy was used to image and compare fluorescent fusions of another carboxysome expression platform pHnCB (with mTurquoise2 fusion) to pXpressome designs. Images captured were used to several various phenotype markers such as cell length **(B)**, cell width **(C)** and punctate level **(D)**. In all cases, sample median is represented by a solid black line and interquartile ranges by dashes black lines. The median of pHnCB-mTQ2 in panels B-D is shown as a dashed gray line for easier comparisons. **** represents p < 0.0001 for pHnCB-mTQ2 and mTagBFP2 samples to all others. Population sizes for these analyses can be found in Supplemental Table S4. Cell health of all designs was evaluated by comparing the growth rates of cells carrying indicated plasmids when either uninduced **(E)** or uninduced **(F)**.

The internal morphology of fluorescent constructs also visually differed between pHnCB and pXpressome samples, with the former being more diffuse and the latter more granular in appearance. We quantified this in terms of a punctate level analysis (Figure 4D). Here, fluorescence distribution was quantified by determining the distribution of normalized pixel intensities per cell. More punctate foci in cells have a higher population of darker pixels, indicating increased clustering into distinct carboxysomes across fewer pixels, while less punctate foci in cells have a more even distribution of pixel intensities. This analysis found that pHnCB-mTQ2 expressing cells have a median punctate level of 0.14 while all pXpressome designs are between 0.18 – 0.45 and indicate more punctate-like carboxysome expression. The lowest punctate level, was given by the mTagBFP2 fusion, was found to be not significantly higher than pHnCB-mTQ2.

We also quantified the effects of these plasmids on cell growth when both uninduced and induced. Uninduced cultures carrying pHnCB tended to grow slower and with more inconsistency than cells carrying pXpressome plasmids (Figure 4E). This is largely attributed to both a longer lag phase and slower overall growth rate. These patterns continued when expression was induced, albeit to a lesser extent (Figure 4F). Meanwhile, pXpressome-carrying cultures all grow consistently, induced or not, regardless of fusion tag. These results make sense given that the primary differences stem from the promoters used and not due to the choice of fusion tags.

## Discussion

BMC research can be facilitated by having robust model systems for study. Plasmid based systems offer a streamlined format that are compatible with higher-throughput expression and manipulation protocols. In this work we introduce a robust plasmid system, the pXpressome toolkit, for researchers to easily adopt carboxysomes into their research. We show that our plasmid system can produce functionally and structurally intact carboxysomes. PCR-based mutagenesis protocols are employed to create and screen several mutants for their structure and activity. Further, a fluorescent carboxysome palette is also developed that can meet any researchers imaging needs. We also demonstrate that our system produces consistent phenotypes and are not inhibiting to cell health.

BMC systems are extremely sensitive to expression parameters. Off-target structures can be common due to a kinetic race between the relative assembly propensities of cargo and/or shell factors which differ between different BMCs.^37,39^ These aberrant assemblies can impact cell health and have been observed interfering with proper septation.^60,61^ Due to this, we thought it was important to test the different promoters used by our system, which uses P_bad_, and that of another carboxysome expression platform pHnCB, which uses P_lacUV5_. While *lacUV5* is stronger than its ancestral *lac* promoter, it can also be leakier which can increase cellular stress and lower protein yields.^62^ The *araBAD* promoter, in contrast, is more tightly regulated and could lead to better outcomes. These general notions were supported by data we collected comparing the two platforms. Cells expressing carboxysomes from P_bad_ on pXpressome plasmids were uniform in their morphology and approximately half the length of those expressing from P_lacuv5_. Cells harboring pXpressome plasmids also grew significantly faster when uninduced. The tag-agnostic morphological and growth consistency shown by the pXpressome system suggests that our choice of a different mTurquoise2 tag in pHnCB is not responsible for the differences observed. These data in total show that carboxysome expression from P_lacuv5_ promoters, and microcompartments at large, may suffer from basal expression which leads to inconsistencies and cell division defects. While not tested here, these negative effects may be abrogated by culturing in the presence of glucose and/or using expression strains which encode for T4 lysozyme as both have been shown to lower basal expression from *lac* promoters. Overall, our data suggests that future synthetic BMC expression schemes may benefit from using highly regulated promoters such as P_bad_.

One surprising result from our simple mutant screen was the size differences in populations of carboxysome lacking either the BMC-P CsoS4A or the BMC-H CsoS1C. Specifically, carboxysomes lacking one copy of BMC-P were larger than wild type and those lacking one copy of the major BMC-H CsoS1C were smaller than wild type (Figure 5). These results can be interpreted within the context of altering the ratios of BMC-H to BMC-P. Computational modeling of BMC assembly have demonstrated that lowering the amount of BMC-P relative to BMC-H in a system can result in larger shells due to exerting less curvature into the system.^63,64^ Our data validates this and suggests that carboxysomes increase in triangulation value from ∼169 (wild type) to ∼225 (ΔCsoS4A).^22^ This also agrees with our observations of elongated and nanocone carboxysomes, which have been speculated to arise due to kinetic traps^38^ during the assembly process when a lack of sufficient BMC-P results in asymmetric pentamer distribution.^36^ Other BMC systems have been observed having elongated/larger structures when all BMC-P are deleted.^35,48^ Conversely, we also show that decreasing the amount of BMC-H relative to BMC-P results in smaller particles with a triangulation value ∼100 (Figure 5). Here, the relatively higher concentration of intracellular BMC-P inflicts more curvature and facets lack the building blocks necessary to extend thereby forming smaller shells. These data offer confirmation of the computational models which show that adapting the shell is a sufficient route to alter carboxysome morphology.^64^ These data also give routes to bias assembly towards elongated or nanocone assemblies that can alter the surface area to volume ratios.

**Figure 5:**
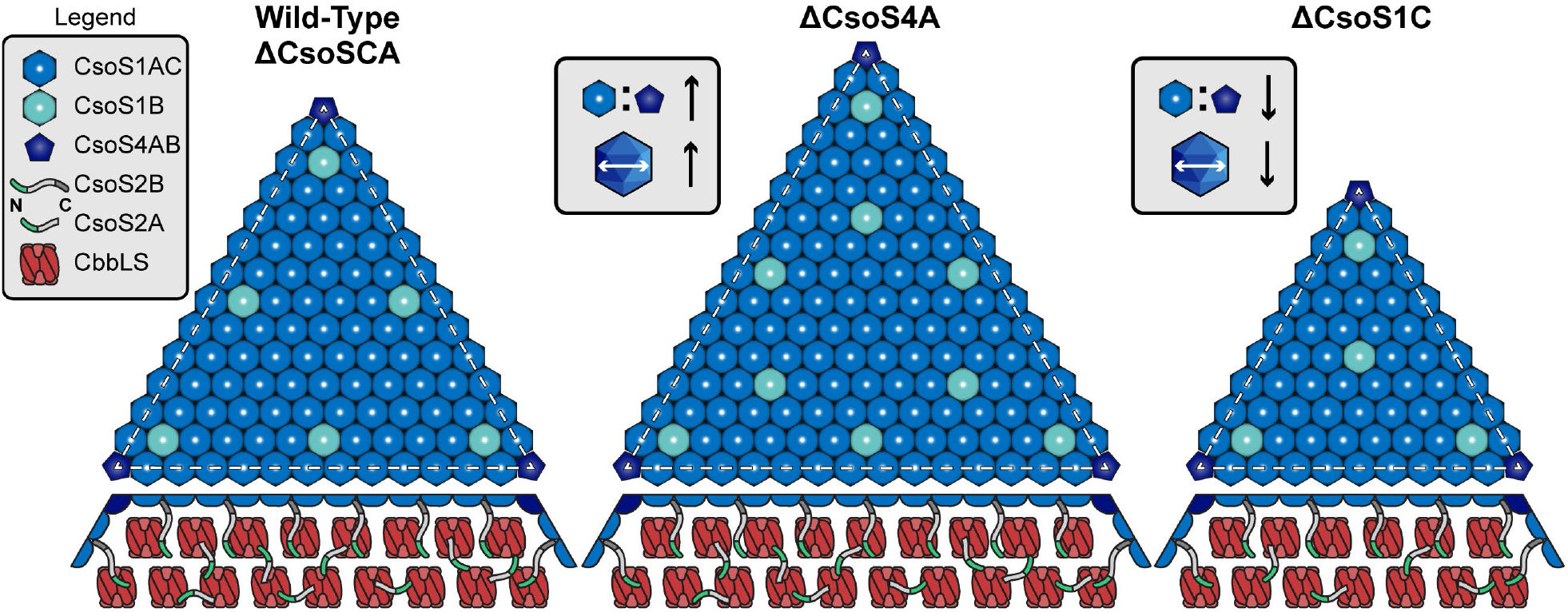
The ratios of structural components influence morphology. Our robust plasmid-based system allowed us to rapidly produce mutant carboxysomes, greatly facilitating analysis. Wild type and carbonic anhydrase (CsoSCA) knockouts are of similar size, as expected, giving rise to a triangulation (T) value of ∼169. In contrast, deleing CsoS4A gave rise to larger carboxysomes with T = ∼225. Lastly, knockout of the major BMC-H CsoS1C leads to smaller particles with T = ∼100. Facet representations shown are anticipated models for this system with CsoS1B placements being speculative and their abundances informed by Sun *et al*. 2022. The latter two mutants demonstrate that the carboxysome is structural plasticity and that the ratios of BMC-H and BMC-P can influence the final morphology.

Other work has shown that altering the domain repeats and the ratios of the two forms of the scaffold CsoS2 can also change carboxysome size by specifically altering the ratio of M-peptide and CTD repeats.^46^ Further, the inclusion of shell-binding C-terminal domain (CTD) repeats has been found to promote the formation of higher diameter synthetic α-carboxysome shells.^19^ These observations and ours suggests that both shell and scaffold-centric avenues may be able to work together synergistically to control carboxysome size. This is supported by our observation that knocking out CsoS1C, leading to smaller shells, require less CsoS2A. We believe that the lower abundance of CsoS2A cannot be explained by the deletion of the CsoS1C reading frame as it is downstream of the CsoS2 gene. One interpretation of this observation is that CsoS2A has its function sufficiently replaced by CsoS2B and that there is a preference, and indeed a requirement,^46,65^ for CsoS2B over CsoS2A. The smaller facets in the ΔCsoS1C carboxysomes, which arise from less available hexameric building blocks, may be sufficiently stabilized by the middle region repeats from CsoS2B.^46,47^ Further, the N-terminal domain of CsoS2B may likewise be able to largely replace the need for CsoS2A for Rubisco networking and inclusion. Overall, our data supports a growing body of evidence that carboxysomes are structurally adaptable to new compositional distributions of structural and scaffold components. These compositions changes can complement reprogramming efforts by using the middle and/or CTD of CsoS2 as proxy “encapsulation peptides” to incorporate new cargoes. ^66,67^

Molecular biology tools are increasingly becoming cheaper and more robust. Cloning methodologies like Gibson assembly coupled to PCR amplification have made concerns about restriction site placement obsolete. Further, the use of custom synthetic DNA fragments to encode desired genes are rapidly becoming cheaper and can accommodate longer (approaching 10 kilobases) and more complex designs which give immense flexibility to ones’ research. It is critical to provide BMC expression formats that are compatible with these ever-evolving tools to increase their accessibility. The pXpressome system fits into this shifting synthetic biology paradigm and lowers the bar to enter BMC research. This toolkit which will enable researchers to redesign carboxysomes for enhanced carbon fixation or serve as a platform for other nanoencapsulation goals.

## Acknowledgements

We would like to thank Dr. Alicia Withrow and the Center for Advanced Microscopy at Michigan State University for performing electron microscopy services. C.R.G.E and D.T. acknowledge funding from by LANL’s Laboratory Direction Research and Development (LDRD) funds (20220387ER). A.V.G acknowledges funding from the National Institute of General Medical Sciences o the National Institutes of Health (Grant no. R25GM152128).

## Author Contributions

**D.S.T.:** conceptualization, formal analysis, investigation, methodology, project administration, visualization, writing – original draft, writing – review and editing. **Y.H.:** formal analysis, investigation, methodology, validation, visualization, writing – original draft, writing – review and editing. **A.G.V.:** supervision, writing – original draft, writing – review and editing. **C.R.G.E.:** funding acquisition, project administration, supervision, writing – original draft, writing – review and editing.

## Materials and Correspondence

All plasmids and associated sequence maps used in this study are available upon request. Please make requests and correspondence out to Dr. Daniel Trettel at.

## Methods

### Molecular Biology and Materials

A complete list of primers, synthetic gene fragments, and plasmid names can be found in the Supplemental Materials. Briefly, all cloning used the NEBuilder® HiFi DNA Assembly Cloning Kit (#E5520S), Q5® High-Fidelity 2X Master Mix (#M0492S) and/or KLD Enzyme Mix (#M0554S) offerings from New England Biolabs. Gibson assemblies were carried out using 0.02 pmol of backbone with a 3-fold molar excess of insert. Reactions were completed at 50°C for 20 minutes and then immediately frozen at -20°C until transformed into NEB5alpha chemically competent *E. coli* (#C2988J). Clone screening was performed on miniprepped plasmid DNA with whole-plasmid sequencing offered by Plasmidsaurus. PCR reactions were performed at 50 µL scale using a 15 s/kb extension time and appropriate primer annealing parameters. PCR reactions were treated with 1 µL of DpnI (#R0176S) to remove template DNA for 30 minutes at 37°C followed by cleanup and concentration using the Zymo DNA Clean and Concentrator kit and quantified with a Qubit dsDNA BR kit. For PCR mutagenesis, 1 µL of successful PCR was mixed with KLD Enzyme Mix according to manufacturer’s protocols prior to transformation.

### Carboxysome Enrichment Protocol

Glycerol stocks of TOP10 strains harboring appropriate plasmid were freshly streaked onto LB plates containing appropriate antibiotic. Single colonies were then used to inoculate 4 mL overnight cultures in 2xYT broth and antibiotic and incubated overnight at 37°C, 250 RPM. Overnight cultures were then used to inoculate 250 – 1500 mL of 2xYT supplemented with antibiotic in baffled flasks and grown to an optical density at 600 nm of ∼0.6-0.8 at 37°C, 250 RPM. At that density, cells were briefly cooled to room temperature prior to the addition of arabinose to 0.02% (w/v) (or ∼1 mM) and continued incubation at 20°C and 250 RPM for 16-18 hours. Cells were harvested the next day via centrifugation (5000 RCF, 10 minutes) and resuspended at 5 mL/g wet cell pellet in B-PERII Bacterial Protein Extraction Reagent (Thermo, #78260) supplemented with 0.5 mM PMSF, 0.5 mg/mL egg white lysozyme and 12.5 U/mL recombinant benzonase (Syd Labs, #BP4200). The suspensions were lysed at 25°C for 40 minutes with constant agitation. Insoluble material was then removed by centrifugation (12000 RCF, 15 minutes, 4°C). The supernatant was decanted into a fresh tube and then centrifuged (30000 RCF, 45 minutes, 4°C) to pellet carboxysomes. The carboxysome pellet was then gently resuspended by pipetting in 0.66x volume of ice-cold TEMB (5 mM Tris-HCl, 1 mM EDTA, 10 mM MgCl_2_, 20 mM NaHCO_3_ pH 8.0). The carboxysomes were then repelleted again (20000 RCF, 30 minutes, 4°C) and resuspended into 3 mL of TEMB, Carry-over cell debris are then removed by several additional rounds of centrifugation (3×12000 RCF, 1 minute, 4°C). Final carboxysome fractions were quantified with a Pierce BCA Assay Kit against a BSA standard. Typical yields were >2 mg/g cell pellet.

### Transmission Electron Microscopy

Carboxysome samples were fixed and diluted from 1 mg/mL stocks prior to deposition onto formvar-carbon coated grids for 5 minutes. Grids were then washed with distilled water, dried and stained with 1% uranyl acetate. Images were taken with a 1400Flash Transmission Electron Microscope (JEOL).

### Dynamic Light Scattering

Light scattering measurements were performed on a Malvern Zetasizer. Samples were diluted to 100 µg/mL in TEMB buffer and equilibrated to 25°C. Triplicate samples were then measured for 3×30s data collections.

### Rubisco Activity Assay

Rubisco activity assays were carried out with a CheKine Micro Rubisco Activity Kit (#MBS9719219) following a modified protocol. The protocol was modified to account for the Rubisco source being enriched carboxysomes as opposed to leaf extracts. Buffers were prepared per the manufacturers protocol and the reaction initiated with the addition of carboxysome samples to a final concentration of 50 µg/mL in a UV-transparent half-area 96-well plate. Changes in absorbance at 340 nm were then measured every 15s for 30 minutes at 30°C in a Biotek Synergy plate reader. Initial velocity was calculated after the initial lag phase from the change in absorbance between 5 to 7 minutes and adjusted for the extinction coefficient of NADH at 340 nm.

### Laser Scanning Confocal Microscopy Imaging

Carboxysome expression was carried out as indicated in the Carboxysome Enrichment Protocol section with samples taken at indicated timepoints. Samples were harvested with centrifugation (3000 RCF, 2 minutes) and resuspended in phosphate buffered saline (PBS) pH 7.4. In the case of gDNA staining, samples in PBS were incubated with SYTO9 for 5 minutes at room temperature prior to repelleting as before and resuspended into an equal volume of PBS. Samples were then spotted into a prepared 1.5% agarose pad in PBS, spread and dried for 5 minutes prior to imaging on an Olympus FV3000. The 405 nm laser line was used to excited mTagBFP2 and mJuniper, 488 nm laser line for mNeonGreen, mChartreuse, and mLemon, and 561 nm laser line for mScarlet-I3 and mLychee. Images were colored in ImageJ and are presented without further modification beyond cropping.

### Wide-Field Fluorescence and Phase-Contrast Imaging

Wide-field fluorescence and phase-contrast imaging were performed using a Nikon Ti2-E motorized inverted microscope controlled by the NIS Elements software with a SOLA 365 LED light source, a 100× objective lens (Oil CFI Plan Apochromat DM Lambda Series for Phase Contrast), and a Photometrics Prime 95B back-illuminated sCMOS camera or Hamamatsu Orca-Flash 4.0 LTS camera. Fusions to mNeonGreen, mJuniper, mLemon, and mChartruese were imaged using a “GFP” filter set (C-FL GFP, Hard Coat, High Signal-to-Noise, Zero Shift, Excitation: 436 ± 10 nm, Emission: 480 ± 20 nm, Dichroic Mirror: 455 nm). Fusions to mTagBFP2 were imaged using a “DAPI” filter set (C-FL DAPI, Hard Coat, High Signal-to-Noise, Zero Shift, Excitation: 350±25 nm, Emission: 460±25 nm, Dichroic Mirror: 400 nm). Fusions to mScarlet-I3 and mLychee were imaged using a “Texas Red” filter set (C-FL Texas Red, Hard Coat, High Signal-to-Noise, Zero Shift, Excitation: 560±20 nm, Emission: 630±37.5 nm; Dichroic Mirror: 585 nm). Cells (2 µl) were then spotted on an imaging agar pad of 1 cm diameter that was made of 1.5% UltraPure agarose in LB media. The cell-containing side of the pad was flipped immediately onto a 35-mm glass bottom dish and mounted onto the stage of the Nikon Ti2-E motorized inverted microscope for image collection.

### Cell Segmentation

Cell segmentation was performed with the Cellpose51 package in Python. To train the model optimally for bacterial cell morphology, 26 raw phase contrast images of cells were manually annotated. To segment the cells using the trained model, a Gaussian blur (standard deviation of Gaussian = 0.066 µm) was applied to the bacterial cell phase-contrast images and the blurred cells were segmented. Cells touching the borders of the image were ignored. Erroneous segmentations were manually corrected using the Cellpose GUI or excluded from further analysis.^68^

### Punctate Level Analysis

The fluorescence intensity for each pixel in a cell was corrected for background by subtracting the median value of all pixels in an image outside of the cell regions and normalized by the minimum and maximum pixel intensity values in the corresponding cell (similar to “condensation coefficients”)^68^:

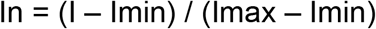

The normalized pixel intensities were then binned to generate histograms that represent the localization pattern for each protein in cells. Homogeneously distributed protein would display a flat distribution while a strongly left-skewed distribution indicates more clustered proteins in the cell. Erroneous foci or cell identifications were excluded from further analysis.

