## Supplemental Information for "A robust synthetic biology toolkit to advance carboxysome study and redesign"

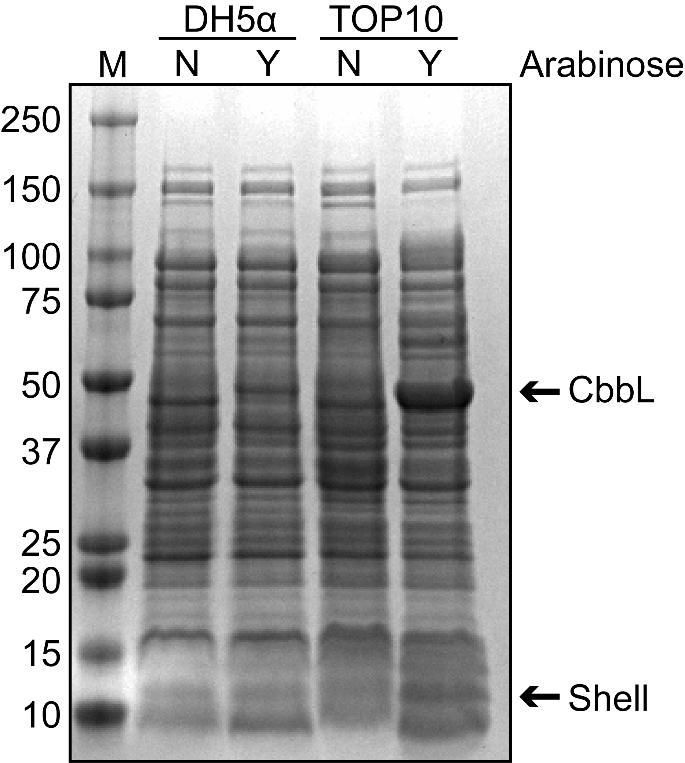

**Figure S1: Comparison of carboxysome expression from pXpressome plasmid in common *E. coli* strains.** Two common lab strains were grown to mid-log then induced (Y) or not (N) at 0.02% (w/v) (~1 mM) arabinose overnight at our standard induction conditions (20°C, 250 RPM). Equivalent volumes of culture were then harvested for SDS-PAGE analysis by pelleting followed by resuspension in 1x SDS-PAGE sample loading buffer and heat treatment at 95°C for 2 minutes. Only the TOP10 cells with added arabinose show the presence of new bands which correspond to carboxysome components.

**Figure S2: The presence of elongated and nanocone carboxysomes in ΔCsoS4A samples.** Various negative stain electron micrographs are shown containing elongated (red arrows) and nanocone (blue arrows) structures. A hybrid structure (purple arrow), appearing as an elongated nanocone, is also observed. The right most micrograph shows a nanocone (top particle) under high magnification.

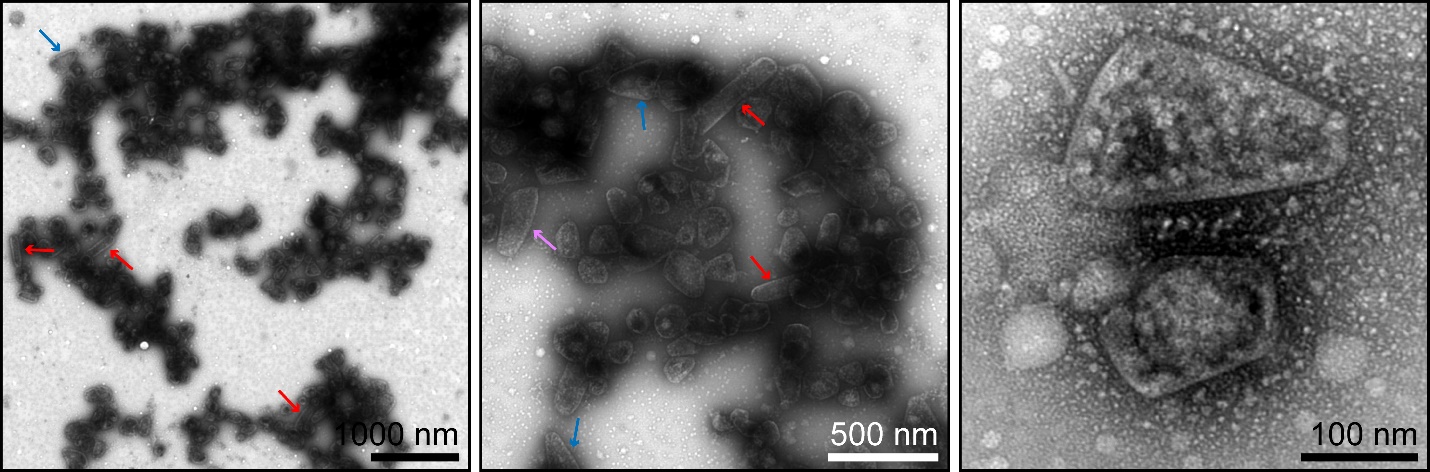

**Table S1: Primers used for various cloning methodologies in this report.**

| **Primer** | **Sequence (5’ to 3’)** | **Purpose** |
| --- | --- | --- |
| oDT097 | CGAATTCGCTAGCCCAAAAAAACG | Linearize pBAD33 |
| oDT098 | ATGAGAGAAGATTTTCAGCCTGATACAG | Linearize pBAD33 |
| oDT115 | TTTTGGGCTAGCGAATTCGatttcagctaggtgcgcg | Amplify CB operon |
| oDT116 | GGCTGAAAATCTTCTCTCATttattagctattcagatttgcgatacaccac | Amplify CB operon |
| oDT126 | cttaatcaaccgcgcgc | Produce ΔCsoSCA |
| oDT127 | gcaggatatcggaagcccg | Produce ΔCsoSCA |
| oDT128 | cgggcttccgatatcctgc | Produce ΔCsoS4A |
| oDT129 | cctctgatttgacgattatcggg | Produce ΔCsoS4A |
| oDT131 | caaaatcaactcatctagcgatgg | Produce ΔCsoS1C |
| oDT132 | ggattgggaaagacgaaccg | Produce ΔCsoS1C |
| oDT139 | gttgccgcggtagacc | FP assemblies |
| oDT140 | gtcaagtctgtcattgcgctg | FP assemblies |
| oDT141 | TTTTTTTAAGGCAGTTATTGGTGCCC | Antibiotic swap |
| oDT142 | ACGTCTCATTTTCGCCAAAAGTTG | Antibiotic swap |

**Table S2: Synthetic gene fragments used for Gibson assembly cloning reactions.** Underlined regions are the assembly overlaps, bolded text is the protein reading frame. All gene fragments were ordered from Twist Biosciences with adapter sequences off and codon optimized for *E. coli* expression. All fragments were resuspended to 40 ng/µL in nuclease-free water and used directly for Gibson assembly.

| **Name** | **Sequence (5’-3’)** | **Purpose** |
| --- | --- | --- |
| gDT046 | CTTTTGGCGAAAATGAGACGTTTTGTTTATTTTTCTAAATACATTCAAATATGTATCCGCTCATGAGACAATAACCCTGATAAATGCTTCAATAATATTGAAAAAGGAAGAGT**ATGAGCCATATTCAACGGGAAACGTCTTGCTCTAGGCCGCGATTAAATTCCAACATGGATGCTGATTTATATGGGTATAAATGGGCTCGCGATAATGTCGGGCAATCAGGTGCGACAATCTATCGATTGTATGGGAAGCCCGATGCGCCAGAGTTGTTTCTGAAACATGGCAAAGGTAGCGTTGCCAATGATGTTACAGATGAGATGGTCAGACTAAACTGGCTGACGGAATTTATGCCTCTTCCGACCATCAAGCATTTTATCCGTACTCCTGATGATGCATGGTTACTCACCACTGCGATCCCCGGGAAAACAGCATTCCAGGTATTAGAAGAATATCCTGATTCAGGTGAAAATATTGTTGATGCGCTGGCAGTGTTCCTGCGCCGGTTGCATTCGATTCCTGTTTGTAATTGTCCTTTTAACAGCGACCGCGTATTTCGTCTCGCTCAGGCGCAATCACGAATGAATAACGGTTTGGTTGATGCGAGTGATTTTGATGACGAGCGTAATGGCTGGCCTGTTGAACAAGTCTGGAAAGAAATGCATAAACTTTTGCCATTCTCACCGGATTCAGTCGTCACTCATGGTGATTTCTCACTTGATAACCTTATTTTTGACGAGGGGAAATTAATAGGTTGTATTGATGTTGGACGAGTCGGAATCGCAGACCGATACCAGGATCTTGCCATCCTATGGAACTGCCTCGGTGAGTTTTCTCCTTCATTACAGAAACGGCTTTTTCAAAAATATGGTATTGATAATCCTGATATGAATAAATTGCAGTTTCATTTGATGCTCGATGAGTTTTTC**TAATTTTTTTAAGGCAGTTATTGGTGCC | KanR |
| gDT047 | CTTTTGGCGAAAATGAGACGTCGCGGAACCCCTATTTGTTTATTTTTCTAAATACATTCAAATATGTATCCGCTCATGAGACAATAACCCTGATAAATGCTTCAATAATATTGAAAAAGGAAGAGT**ATGAGTATTCAACATTTCCGTGTCGCCCTTATTCCCTTTTTTGCGGCATTTTGCCTTCCTGTTTTTGCTCACCCAGAAACGCTGGTGAAAGTAAAAGATGCTGAAGATCAGTTGGGTGCACGAGTGGGTTACATCGAACTGGATCTCAACAGCGGTAAGATCCTTGAGAGTTTTCGCCCCGAAGAACGTTTTCCAATGATGAGCACTTTTAAAGTTCTGCTATGTGGCGCGGTATTATCCCGTGTTGACGCCGGGCAAGAGCAACTCGGTCGCCGCATACACTATTCTCAGAATGACTTGGTTGAGTACTCACCAGTCACAGAAAAGCATCTTACGGATGGCATGACAGTAAGAGAATTATGCAGTGCTGCCATAACCATGAGTGATAACACTGCGGCCAACTTACTTCTGACAACGATCGGAGGACCGAAGGAGCTAACCGCTTTTTTGCACAACATGGGGGATCATGTAACTCGCCTTGATCGTTGGGAACCGGAGCTGAATGAAGCCATACCAAACGACGAGCGTGACACCACGATGCCTGCAGCAATGGCAACAACGTTGCGCAAACTATTAACTGGCGAACTACTTACTCTAGCTTCCCGGCAACAATTAATAGACTGGATGGAGGCGGATAAAGTTGCAGGACCACTTCTGCGCTCGGCCCTTCCGGCTGGCTGGTTTATTGCTGATAAATCTGGAGCCGGTGAGCGTGGGTCTCGCGGTATCATTGCAGCACTGGGGCCAGATGGTAAGCCCTCCCGTATCGTAGTTATCTACACGACGGGGAGTCAGGCAACTATGGATGAACGAAATAGACAGATCGCTGAGATAGGTGCCTCACTGATTAAGCATTGGTAA**TTTTTTTAAGGCAGTTATTGGTGCC | AmpR |
| gDT048 | ttcgtggtctaccgcggcaacGGCAGC**GTGAGCAAAGGCGAAGAAGATAACATGGCGAGCCTGCCGGCGACCCATGAACTGCATATTTTTGGCAGCATTAACGGCGTGGATTTTGATATGGTGGGCCAGGGCACCGGCAACCCGAACGATGGCTATGAAGAACTGAACCTGAAAAGCACCAAAGGCGATCTGCAGTTTAGCCCGTGGATTCTGGTGCCGCATATTGGCTATGGCTTTCATCAGTATCTGCCGTATCCGGATGGCATGAGCCCGTTTCAGGCGGCGATGGTGGATGGCAGCGGCTATCAGGTGCATCGCACCATGCAGTTTGAAGATGGCGCGAGCCTGACCGTGAACTATCGCTATACCTATGAAGGCAGCCATATTAAAGGCGAAGCGCAGGTGAAAGGCACCGGCTTTCCGGCGGATGGCCCGGTGATGACCAACAGCCTGACCGCGGCGGATTGGTGCCGCAGCAAAAAAACCTATCCGAACGATAAAACCATTATTAGCACCTTTAAATGGAGCTATACCACCGGCAACGGCAAACGCTATCGCAGCACCGCGCGCACCACCTATACCTTTGCGAAACCGATGGCGGCGAACTATCTGAAAAACCAGCCGATGTATGTGTTTCGCAAGACCGAACTGAAACATAGCAAAACGGAGCTGAACTTTAAAGAATGGCAGAAAGCGTTTACCGATGTGATGGGCATGGATGAACTGTATAAA**TAAgtcaagtctgtcattgcgct | mNeonGreen |
| gDT049 | ttcgtggtctaccgcggcaacGGCAGC**GATAGCACCGAAGCGGTGATTAAAGAATTTATGCGCTTTAAAGTGCATATGGAAGGCAGCATGAACGGCCATGAATTTGAAATTGAAGGCGAAGGCGAAGGCCGCCCGTATGAAGGCACCCAGACCGCGAAACTGAAAGTGACCAAAGGCGGCCCGCTGCCGTTTAGCTGGGATATTCTGAGCCCGCAGTTTATGTATGGCAGCCGCGCGTTTATTAAACATCCGGCGGATATTCCGGATTATTGGAAACAGAGCTTTCCGGAAGGCTTTAAATGGGAACGCGTGATGATTTTTGAAGATGGCGGCACCGTGAGCGTGACCCAGGATACCAGCCTGGAAGATGGCACCCTGATTTATAAAGTGAAACTGCGCGGCGGCAACTTTCCGCCGGATGGCCCGGTGATGCAGAAACGCACCATGGGCTGGGAAGCGAGCACCGAACGCCTGTATCCGGAAGATGTGGTGCTGAAAGGCGATATTAAAATGGCGCTGCGCCTGAAAGATGGCGGCCGCTATCTGGCGGATTTTAAAACCACCTATAAAGCGAAAAAACCGGTGCAGATGCCGGGCGCGTTTAACATTGATCGCAAACTGGATATTACCAGCCATAACGAAGATTATACCGTGGTGGAACAGTATGAACGCAGCGTGGCGCGCCATAGCACCGGCGGCAGCGGCGGCAGC**TAAgtcaagtctgtcattgcgct | mScarlet-I3 |
| gDT050 | ttcgtggtctaccgcggcaacGGCAGC**GTGAGCAAAGGCGAAGAACTGATTAAAGAAAACATGCATATGAAACTGTATATGGAAGGCACCGTGGATAACCATCATTTTAAATGCACCAGCGAAGGCGAAGGCAAACCGTATGAAGGCACCCAGACCATGCGCATTAAAGTGGTGGAAGGCGGCCCGCTGCCGTTTGCGTTTGATATTCTGGCGACCAGCTTTCTGTATGGCAGCAAAACCTTTATTAACCATACCCAGGGCATTCCGGATTTTTTTAAACAGAGCTTTCCGGAAGGCTTTACCTGGGAACGCGTGACCACCTATGAAGATGGCGGCGTGCTGACCGCGACCCAGGATACCAGCCTGCAGGATGGCTGCCTGATTTATAACGTGAAAATTCGCGGCGTGAACTTTACCAGCAACGGCCCGGTGATGCAGAAAAAAACCCTGGGCTGGGAAGCGTTTACCGAAACCCTGTATCCGGCGGATGGCGGCCTGGAAGGCCGCAACGATATGGCGCTGAAACTGGTGGGCGGCAGCCATCTGATTGCGAACGCGAAAACCACCTATCGCAGCAAAAAACCGGCGAAAAACCTGAAAATGCCGGGCGTGTATTATGTGGATTATCGCCTGGAACGCATTAAAGAAGCGAACAACGAAACCTATGTGGAACAGCATGAAGTGGCGGTGGCGCGCTATTGCGATCTGCCGAGCAAACTGGGCCATAAACTGAAC**TAAgtcaagtctgtcattgcgct | mTagBFP2 |
| gDT051 | ttcgtggtctaccgcggcaacGGCAGC**GATAGCACCGAAGCGATTATTAAAGAATTTATGCGCTTTAAAGTGCATATGGAAGGCAGCGTGAACGGCCATGAATTTGAAATTGAAGGCGAAGGCGAAGGCCGCCCGTATGAAGCGTTTCAGACCGCGAAACTGAAAGTGACCAAAGGCGGCCCGCTGCCGTTTGCGTGGGATATTCTGAGCCCGCAGTTTATGTATGGCAGCAAAGCGTATATTAAACATCCGGCGGATATTCCGGATTATTTTAAACAGAGCTTTCCGGAAGGCTTTCGCTGGGAACGCGTGATGAACTTTGAAGATGGCGGCATTATTCATGTGAACCAGGATAGCAGCCTGCAGGATGGCGTGTTTATTTATAAAGTGAAACTGCGCGGCACCAACTTTCCGCCGGATGGCCCGGTGATGCAGAAACGCACCATGGGCTGGGAACCGAGCGAAGAACGCATGTATCCGGAAGATGGCGCGCTGAAAAGCGAAATTAAAAAACGCCTGAAACTGAAAGATGGCGGCCATTATGCGGCGGAAGTGAAAACCACCTATAAAGCGAAAAAACCGGTGCAGCTGCCGGGCGCGTATATTGTGGATATTAAACTGGATATTGTGAGCCATAACGAAGATTATACCGTGGTGGAACAGTATGAACGCGCGGAAGGCCGCCATAGCGGCAGCCAGGGCGGCAGCGGCGGCAGCCTGTATAAA**TAAgtcaagtctgtcattgcgct | mLychee |
| gDT052 | ttcgtggtctaccgcggcaacGGCAGC**AGCAAAGGCGAAGAACTGTTTACCGGCGTGGTGCCGATTCTGGTGGAACTGGATGGCGATGTGAACGGCCATAAATTTAGCGTGCGCGGCGAAGGCGAAGGCGATGCGACCATTGGCAAACTGACCCTGAAATTTATTTGCACCACCGGCAAACTGCCGGTGCCGTGGCCGACCCTGGTGACCACCCTGACCTGGGGCGTGCAGTGCTTTGCGCGCTATCCGGATCATATGAAACAGCATGATTTTTTTAAAAGCGCGATGCCGGAAGGCTATGTGCAGGAACGCACCATTAGCTTTAAAGATGATGGCACCTATAAAACCCGCGCGGAAGTGAAATTTGAAGGCGATACCCTGGTGAACCGCATTGAACTGAAAGGCAGCGGCTTTAAAGAAGATGGCAACATTCTGGGCCATAAACTGGAATATAACTATTTTAGCGATAAAGTGTATATTACCGCGGATAAACAGAAAAACGGCATTAAAGCGAACTTTAAAATTCGCCATAACGTGGAAGATGGCAGCGTGCAGCTGGCGGATCATTATCAGCAGAACACCCCGATTGGCGATGGCCCGGTGCTGCTGCCGGATAACCATTATCTGAGCACCCAGAGCAAACTGAGCAAAGATCCGAACGAAAAACGCGATCATATGGTGCTGCTGGAATTTGTGACCGCGGCGGGCATTACCCATGGCATGGATGAACTGTATAAA**TAAgtcaagtctgtcattgcgct | mJuniper |
| gDT053 | ttcgtggtctaccgcggcaacGGCAGC**AGCAAAGGCGAAGAACTGTTTACCGGCGTGGTGCCGATTCTGGTGGAACTGGATGGCGATGTGAACGGCCATAAATTTAGCGTGCGCGGCGAAGGCGAAGGCGATGCGACCATTGGCAAACTGACCCTGAAATTTATTTGCACCACCGGCAAACTGCCGGTGCCGTGGCCGACCCTGGTGACCAGCCTGGGCTATGGCGTGCAGTGCTTTGCGCGCTATCCGGATCATATGAAACAGCATGATTTTTTTAAAAGCGCGATGCCGGAAGGCTATGTGCAGGAACGCACCATTAGCTTTAAAGATGATGGCACCTATAAAACCCGCGCGGAAGTGAAATTTGAAGGCGATACCCTGGTGAACCGCATTGAACTGAAAGGCAGCGGCTTTAAAGAAGATGGCAACATTCTGGGCCATAAACTGGAATATAACTATAACAGCCATAAAGTGTATATTACCGCGGATAAACAGAAAAACGGCATTAAAGCGAACTTTAAAATTCGCCATAACGTGGAAGATGGCAGCGTGCAGCTGGCGGATCATTATCAGCAGAACACCCCGATTGGCGATGGCCCGGTGCTGCTGCCGGATAACCATTATCTGAGCTATCAGAGCAAACTGAGCAAAGATCCGAACGAAAAACGCGATCATATGGTGCTGCTGGAATTTCTGACCGCGGCGGGCATTACCCATGGCATGGATGAACTGTATAAA**TAAgtcaagtctgtcattgcgct | mLemon |
| gDT054 | ttcgtggtctaccgcggcaacGGCAGC**AGCAAAGGCGAAGAACTGTTTACCGGCGTGGTGCCGATTCTGGTGGAACTGGATGGCGATGTGAACGGCCATAAATTTAGCGTGCGCGGCGAAGGCGAAGGCGATGCGACCATTGGCAAACTGACCCTGAAATTTATTTGCACCACCGGCAAACTGCCGGTGCCGTGGCCGACCCTGGTGACCACCCTGACCTATGGCGTGCAGTGCTTTAGCCGCTATCCGGATCATATGAAACAGCATGATTTTTTTAAAAGCGCGATGCCGGAAGGCTATGTGCAGGAACGCACCATTAGCTTTAAAGATGATGGCACCTATAAAACCCGCGCGGAAGTGAAATTTGAAGGCGATACCCTGGTGAACCGCATTGAACTGAAAGGCAGCGGCTTTAAAGAAGATGGCAACATTCTGGGCCATAAACTGGAATATAACTATAACAGCCATAAAGTGTATATTACCGCGGATAAACAGAAAAACGGCATTAAAGCGAACTTTAAAATTCGCCATAACGTGGAAGATGGCAGCGTGCAGCTGGCGGATCATTATCAGCAGAACACCCCGATTGGCGATGGCCCGGTGCTGCTGCCGGATAACCATTATCTGAGCACCCAGAGCAAACTGAGCAAAGATCCGAACGAAAAACGCGATCATATGGTGCTGCTGGAATTTGTGACCGCGGCGGGCATTACCCATGGCATGGATGAACTGTATAAA**TAAgtcaagtctgtcattgcgct | mChartreuse |

**Table S3: List of all plasmids developed and used in this study**

| **Name** | **In-House Name** | **Description** |
| --- | --- | --- |
| pXpressome1 | pDT062 | Expresses Hneap alpha-carboxysome |
| pXpressome2 | pDT072 | Antibiotic cassette swapped from CmR to KanR |
| pXpressome3 | pDT073 | Antibiotic cassette swapped from CmR to AmpR |
| pXpressome1_cbbS-mNeonGreen | pDT074 | cbbS C-terminally fused to mNeonGreen |
| pXpressome1_cbbS-mScarletI3 | pDT075 | cbbS C-terminally fused to mScarlet-I3 |
| pXpressome1_cbbS-mTagBFP2 | pDT076 | cbbS C-terminally fused to mTagBFP2 |
| pXpressome1_cbbS-mLychee | pDT077 | cbbS C-terminally fused to mLychee |
| pXpressome1_cbbS-mJuniper | pDT078 | cbbS C-terminally fused to mJuniper |
| pXpressome1_cbbS-mLemon | pDT079 | cbbS C-terminally fused to mLemon |
| pXpressome1_cbbS-Chartreuse | pDT080 | cbbS C-terminally fused to mChartreuse |
| pXpressome1_ΔcsoSCA | pDT087 | csoSCA deleted by PCR mutagenesis |
| pXpressome1_ΔcsoS4A | pDT088 | csoS4A deleted by PCR mutagenesis |
| pXpressome1_ΔcsoS1C | pDT089 | csoS1C deleted by PCR mutagenesis |

**Table S4: Number of cells analyzed for Figure 4**

|  | **pHnCB-mTQ2** | **pDT074** | **pDT075** | **pDT076** | **pDT077** | **pDT078** | **pDT079** | **pDT080** |
| --- | --- | --- | --- | --- | --- | --- | --- | --- |
| Cell length | 1653 | 1377 | 524 | 2102 | 1095 | 1082 | 2307 | 2215 |
| Cell width | 1653 | 1377 | 524 | 2102 | 1095 | 1082 | 2307 | 2215 |
| Punctate level | 1087 | 600 | 524 | 600 | 600 | 600 | 600 | 500 |
